# Abnormal brain state distribution and network connectivity in a *SYNGAP1* rat model

**DOI:** 10.1101/2022.02.04.479013

**Authors:** Ingrid Buller-Peralta, Jorge Maicas-Royo, Zhuoen Lu, Sally M. Till, Emma R. Wood, Peter C. Kind, Javier Escudero, Alfredo Gonzalez-Sulser

**Author notes:** **Corresponding author:** Alfredo Gonzalez-Sulser, Simons Initiative for the Developing Brain Fellow, University of Edinburgh, Centre for Discovery Brain Sciences, 1 George Square, Edinburgh, EH89JZ, United Kingdom. Equal contribution (Ingrid Buller-Peralta, Jorge Maicas-Royo, Zhuoen Lu).

## Abstract

Mutations in the *SYNGAP1* gene are one of the common predictors of neurodevelopmental disorders, commonly resulting in individuals developing autism, intellectual disability, epilepsy, and sleep deficits. EEG recordings in neurodevelopmental disorders show potential to identify clinically translatable biomarkers to both diagnose and track the progress of novel therapeutic strategies, as well as providing insight into underlying pathological mechanisms. In a rat model of *SYNGAP1* haploinsufficiency in which the exons encoding the calcium/lipid binding and GTPase activating protein (GAP) domains have been deleted (*Syngap*^+/*Δ-GAP*^) we analysed the duration and occurrence of wake, non rapid eye movement (NREM) and rapid eye movement (REM) brain states during 6 hour multi-electrode EEG recordings. We find that although *Syngap*^+/*Δ-GAP*^ animals spend an equivalent percent time in wake and sleep states, they have an abnormal brain state distribution as the number of wake and NREM bouts are reduced and there is an increase in the average duration of both wake and NREM epochs. We perform connectivity analysis by calculating the average imaginary coherence between electrode pairs at varying distance thresholds during these states. In group averages from pairs of electrodes at short distances from each other, a clear reduction in connectivity during NREM is present between 11.5 Hz and 29.5 Hz, a frequency range that overlaps with sleep spindles, oscillatory phenomena thought to be important for normal brain function and memory consolidation. Sleep spindles occurrence, amplitude, power and spread across multiple electrodes were not reduced in *Syngap*^+/*Δ-GAP*^ rats, with only a small decrease in duration detected. Nonetheless, by analysing the dynamic imaginary coherence during sleep spindles, we found a reduction in high connectivity instances between short-distance electrode pairs. Finally, by comparing the dynamic imaginary coherence during sleep spindles between individual electrode pairs, we identified a group of channels over the right somatosensory, association and visual cortices that have a significant reduction in connectivity during sleep spindles in mutant animals. These data suggest that *Syngap*^+/*Δ-GAP*^ rats have altered brain state dynamics and EEG connectivity, which may have clinical relevance for *SYNGAP* haploinsufficiency in humans.

## Introduction

Neurodevelopmental disorders (NDDs) encompass multiple disease phenotypes that are linked to a highly heterogenous genetic architecture^1^. Mutations in the *SYNGAP1* gene account for as many as 1% of NDD cases, with patients often presenting with intellectual disability (ID) and autism spectrum disorder^2–6^. Most patients with *SYNGAP1* pathogenic variants display some form of epilepsy with a high prevalence of absence seizures and a behavioural developmental delay occurring upon seizure onset. Therefore, *SYNGAP1* haploinsufficiency is often classed as an epileptic encephalopathy^3,7,8^. Additionally, sleep impairments have been reported by parents ^3^ and have been documented in the clinical setting in a high proportion of cases^8,9^.

Characterization of novel biomarkers of NDDs may allow for early diagnosis and rapid therapeutic intervention, enabling quantitative monitoring of treatment efficacy, which is likely critical for the success of future clinical trials^10^. EEG recordings are a potentially reliable method of identifying biomarkers as they provide a fast and direct measure of overall brain activity^11^. However, given the heterogeneity of aetiologies, phenotypes and disease trajectories of NDDs, biomarkers specific for particular genetic disorders such as *SYNGAP1* are imperative to differentiate between NDD types and identify likely disease outcomes^12^.

The rare incidence of NDD patients with specific mutations makes performing large-cohort EEG studies to identify biomarkers difficult. *SYNGAP1* haploinsufficiency cases are thought to occur in only 6.1 per 100,000 people^13^. Identifying clinically relevant biomarkers in genetically modified rodent models of NDDs is a plausible alternative strategy.

*SYNGAP1* encodes a Ras-GTPase-activating protein, which is mainly expressed in the synapses of excitatory neurons^14,15^. It is a key regulator of the postsynaptic density and in synaptic development and plasticity^16^. We recently reported on a novel rat model of *SYNGAP1* haploinsufficiency in which the calcium/lipid binding (C2) and GTPase activating (GAP) domains, areas thought to be critical for the normal function of SYNGAP, have been deleted^17^. Animals heterozygous for the C2/GAP domain deletion (*Syngap*^+/*Δ-GAP*^) displayed reduced exploration and fear extinction, altered social behaviour and spontaneous absence seizures. SYNGAP plays a critical role in synaptic transmission and organisation, therefore we hypothesize that *Syngap*^+/*Δ-GAP*^ animals have abnormal functional connectivity between different cortical regions during specific brain states.

To record seizures, we previously performed 6-hr recordings utilizing skull-surface 32-channel EEG grids, as these mimic human high-density EEG recordings^18^. Here, we perform detailed analysis of the occurrence of multiple brain states during those recordings and find abnormalities in *Syngap*^+/*Δ-GAP*^ rats in the duration and number of sleep and wake occurrences during recordings, as well as differences in connectivity between cortical regions that may have value as clinically translatable biomarkers.

## Materials and Methods

### Animals

This paper uses data gathered during experiments for which some results have been previously published^17^. All animal procedures were undertaken in accordance with the University of Edinburgh animal welfare committee regulations and were performed under a UK Home Office project license. Long Evans-SG^em2/PWC^, hereafter referred to as *Syngap*^+/*Δ-GAP*^ were kept on a 12h/12h light dark cycle with ad libitum access to water and food. Animals were genotyped by PCR. 12 *Syngap*^+/*Δ-GAP*^ and 12 *Syngap*^+/*+*^ animals were recorded and used across all analyses.

### Surgery

15 to 16 week-old *Syngap*^+/*Δ-GAP*^ and *Syngap*^+/*+*^ male rates were anaesthetised with isoflurane and mounted on a stereotaxic frame. Two craniotomies were drilled for bilateral anchor screw placement (+4.0 mm AP, ± 0.5 mm ML) and one for ground screw implantation (−11.5 mm AP, 0.5 mm ML) (A2 Din M1×3 cheese head screw, Screwsandmore, UK), according to the frontal and caudal edges of the EEG array probe (H32-EEG – NeuroNexus, USA). The EEG probe was placed on the skull with its cross-symbol reference point aligned over bregma (Supplementary Fig. 1). The ground electrode and screw were connected with silver paint, and the implant was covered with dental cement. Animals were allowed to recover for a minimum of one-week post-surgery and were housed individually after surgery to prevent damage to the implant. Animals were monitored for any welfare issues arising during or after surgery as well as changes in behaviour, such as less food consumption or decreased responses to stimuli in cages, none were found.

### EEG recordings

Prior to recording, rats were habituated for 20 to 30 minutes to the room. On recording days, up to 4 rats, were placed in individual side-by-side cages inside a 1 × 1 m faraday enclosure. Experimenters were blind to genotype. 6 hr EEG recordings, starting at zeitgeber time (ZT) 3 to 9 (under a 12 light hr: 12 dark hr schedule starting at 07:00 am) were acquired with an OpenEphys acquisition system (OpenEphys, Portugal), through individual 32-channel recording headstage amplifiers with accelerometers (RHD2132 Intantech, USA), at a sampling rate of 1 kHz.

### Visual sleep scoring and absence seizure detection

Off-line visual brain state scoring blind to animal genotype was performed, assigning 5 s epochs to non-rapid eye movement sleep (NREM), rapid eye movement sleep (REM) or wake. NREM epochs displayed high-amplitude slow wave (1 - 4 Hz) EEG activity accompanied by sleep spindles (12 - 17 Hz) and decreased accelerometer activity. REM was identified by sustained theta (5 - 10 Hz) and no accelerometer activity. Wake was identified by the presence of desynchronized EEG and varying levels of accelerometer activity. Although no further analysis on absence seizures is presented here we previously quantified and reported their occurrence^17^ and excluded their occurrence times from our brain state analyses. The electrographical correlate of absence seizures, spike and wave discharges (SWDs), were identified visually and then analysis was confirmed with an automated absence seizure detection algorithm. Briefly, SWDs are characterized by periodic high amplitude oscillations in the theta band between 5 and 10 Hz^19^ which correlates with a spontaneous stop in animal movement. Spectral analysis was performed that identified harmonic peaks in the power spectral density. The code used for analysis is available at https://github.com/Gonzalez-Sulser-Team/SWD-Automatic-Identification^17^.

### Automatic detection of sleep spindles

Automatic detection of sleep spindles during visually scored NREM sleep epochs was performed using the Matlab “Sleepwalker” toolbox with the “sw_run_delta_LFP.m” function (https://gitlab.com/ubartsch/sleepwalker)^20^. EEG signal from electrode placed over S1-Tr right were band-passed filtered between 12 - 17 Hz, with a minimum-maximum length between 0.2 - 3 s, a minimal time gap between events of 0.2 s, amplitude between 25 - 750 μV, a start to end limit threshold of 1.5 SD with a detection limit of 3 SD of the envelope and noise exclusion at ≥ 35 SD.

To detect spindles detected in multiple sites during NREM periods from electrodes that were less than 2 mm away from each other, we used the Python “Yasa” library function “spindles_detect” (https://zenodo.org/record/4632409) to identify spindles with a frequency range of 12 -17 Hz. The comparison was made specifically in electrodes that were within 2 mm distances from each other.

### Spectral power and imaginary coherence analysis

Movement artefacts, identified visually as data points with values greater than 750 µV were discarded from further analysis. Spectral power for each individual epoch for all brain states (wake, NREM and REM) was calculated from an individual channel over the right primary somatosensory as the mean log_10_ power in the 0.2 - 48 Hz range (0.2 Hz steps) using the Multitaper package (Rahim, Burr & Thomson., 2014^5^) for R-studio (RStudio Team, 2020^6^). Epochs were then averaged per brain states for each individual animal.

For imaginary coherence averages across electrode pairs, the data were downsampled by a factor of 8 with the Python “Scipy.signal” function “decimate”, which includes an order 8 Chebyshev type I filter for antialiasing, to increase processing speed. We calculated the coherence for each of the 32 pairs of electrodes in individual brains states in 99 frequency bins (1 – 50 Hz, 0.5 Hz bin size) with the Python Scipy “signal” function “coherence” and extracted the imaginary component using the Python Numpy function “imag”^21^. To ensure statistical normality, coherence values (*R*^*2*^) from each 0.5 Hz frequency were *z*-transformed using Fisher’s *r* to *z, z*-scores were then averaged at every 0.5 Hz bin and re-transformed utilizing the Fisher inverse function to obtain the *Z*^*-1*^ coherence value per electrode pair and frequency band^22^.

We calculated the Euclidean distance between all pairs of electrode using the Python SciPy “Spatial.distance” function “pdist”, with 1.3 mm being the shortest distance between adjacent electrodes and 13.9 mm the longest distance electrode leads. We then utilized varying distance thresholds (from 2 - 10 mm in 1 mm increments) to compare imaginary coherence averages from short and long distances electrode pairs, between the groups of *Syngap*^+/*Δ-GAP*^ and *Syngap*^+/*+*^ animals (Fig. 2A, Fig. 2D, Fig. 3G, Fig. 3A, Fig. 3D, Fig. 3G, and Supplementary Fig. 1 and Supplementary Fig. 2)^22^.

**Figure 1.**
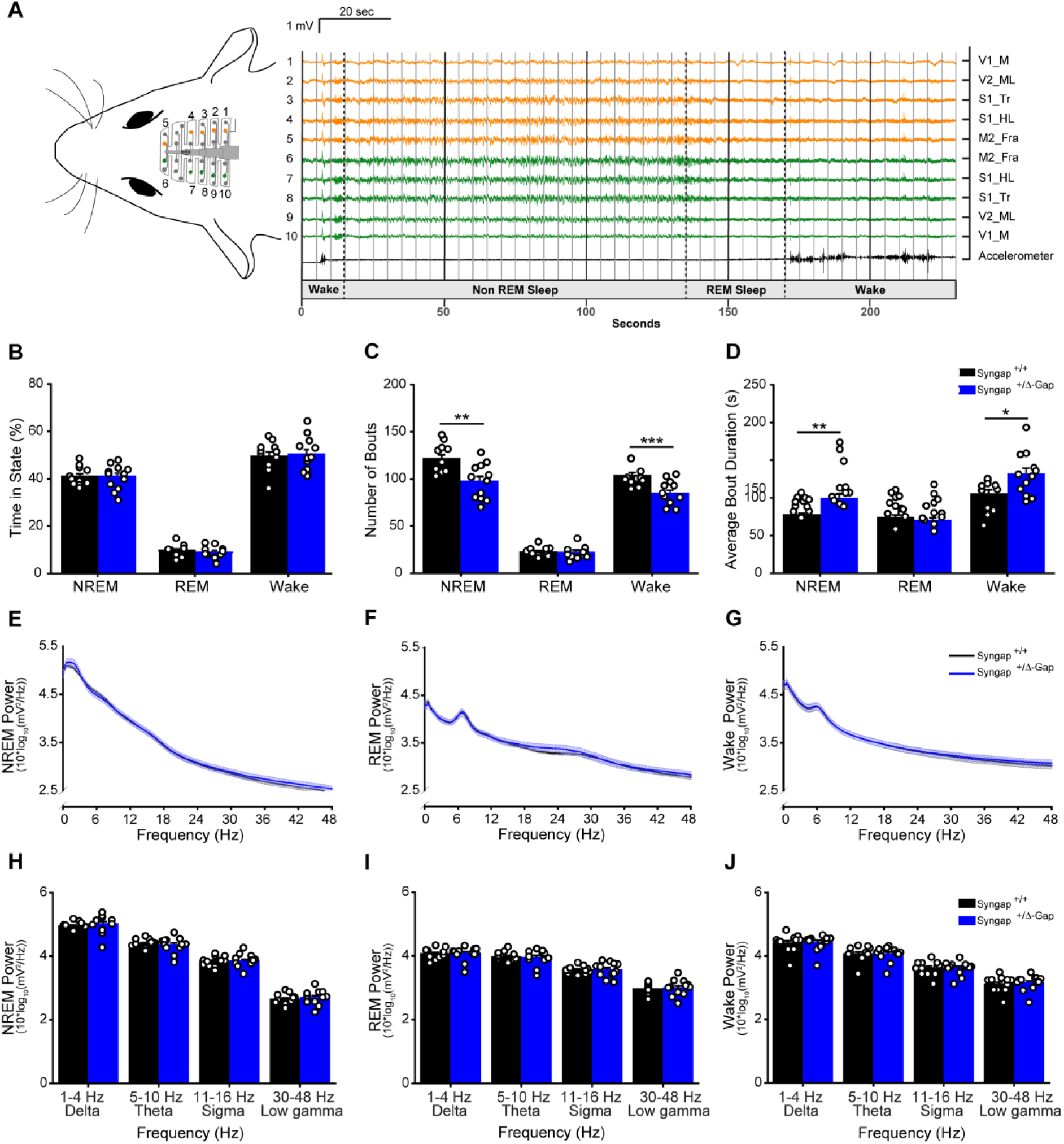
Brain state abnormalities in Syngap^+/Δ-GAP^ rats. (A) Schematic of a 32-channel skull-surface EEG implant illustrating approximate location of electrodes relative to the brain (left). Representative EEG voltage and accelerometer traces from numbered electrodes in schematic on left showing examples from NREM, REM and wake states (right). Dotted black lines indicate brain state transitions, grey lines show example 5 sec brain state epochs. Electrode position abbreviations (Supplementary Fig. 1) are displayed to the right of traces. Plots of percent time of 6 hr recording (B), number of bouts (C) and average bout duration (D) in NREM, REM and wake brain states. Bars indicate mean values (mean ± standard error of the mean (SEM)). Points correspond to values from individual rats. Number of bouts was significantly lower, while bout duration was significantly higher in Syngap^+/Δ-GAP^ rats (* = p<0.05, ** = p<0.01, *** p = <0.001, **** p = <0.0001, unpaired two-sample t-tests and two-sample Man Whitney-U rank sum tests specified in results text). Power spectrum estimates averaged across all NREM (E), REM (F) and wake (G) epochs. Error bars indicate SEM. Plots of average power in commonly used frequency bands during NREM (H), REM (I) and wake (J). Bars indicate mean values (mean ± SEM). Points correspond to values from individual rats. There were no significant differences across any bands between genotypes (unpaired two-sample t-tests and two-sample Man Whitney-U rank sum tests specified in results text).

**Figure 2.**
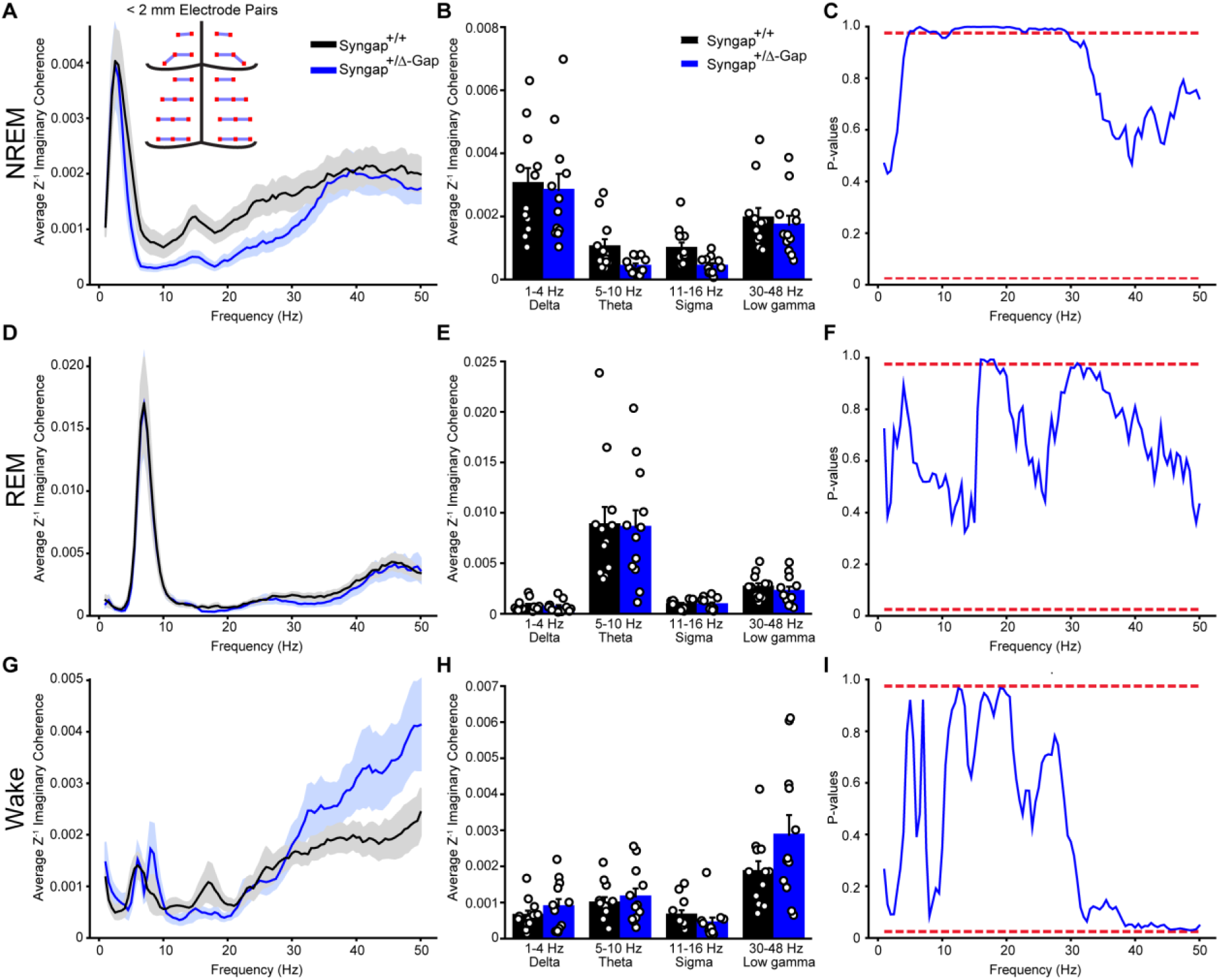
Decreased imaginary coherence during NREM in short-distance (< 2 mm apart) electrode pairs in Syngap^+/Δ-GAP^ rats. Average Z’ imaginary coherence during NREM (A), REM (D) and wake (G) epochs. Shaded area indicates SEM. Inset in A: schematic of electrode pairs < 2mm apart. Plots of average Z’ imaginary coherence in commonly used frequency bands during NREM (B), REM (E) and wake (H). Bars indicate mean values (mean ± SEM). Points correspond to values from individual rats. There were no significant differences across any commonly used bands between genotypes (two-way ANOVA, p = > 0.05). Plots of p-values for cluster-based nonparametric tests during NREM (C), REM (F) and wake (I). Dotted red lines indicate two-sided p-value thresholds of ≥ 0.975 and ≤ 0.025 corresponding to significantly different thresholds equivalent to p ≤ 0.05, two sided. Note: A long cluster of significant frequencies was found during NREM between 11.5 and 29.5 Hz indicating a decrease in Z’ imaginary coherence in Syngap^+/Δ-GAP^ rats.

**Figure 3.**
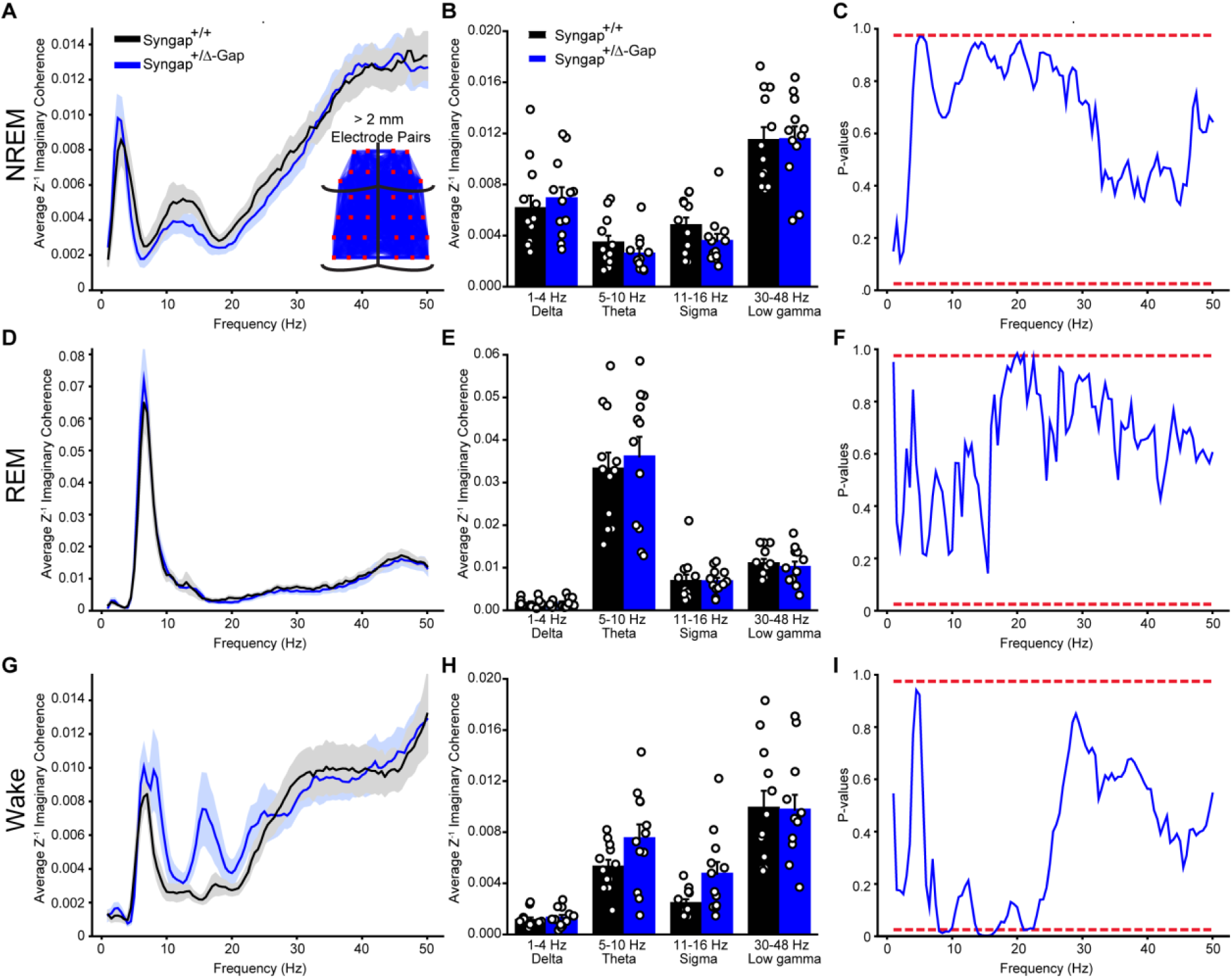
Increased imaginary coherence during wake in long-distance (> 2 mm apart) electrode pairs in Syngap^+/Δ-GAP^ rats. Average Z’ imaginary coherence during REM (A), NREM (D) and wake (G) epochs. Shaded area indicates SEM. Inset in A: schematic of electrode pairs > 2mm apart. Plots of average Z’ imaginary coherence in commonly used frequency bands during NREM (B), REM (E) and wake (H). Bars indicate mean values (mean ± SEM). Points correspond to values from individual rats. There were no significant differences across any commonly used bands between genotypes (two-way ANOVA, p = > 0.05). Plots of p-values for cluster-based nonparametric test during NREM (C), REM (F) and wake (I). Dotted red lines indicate two-sided p-value thresholds of ≥ 0.975 and ≤ 0.025 corresponding to significantly different thresholds equivalent to p ≤ 0.05, two sided. Note: Clusters of significant frequencies were found during wake between 8 and 9.5 Hz and 14 and 16.5 Hz indicating an increase in Z’ imaginary coherence in Syngap^+/Δ-GAP^ rats.

Imaginary phase coherogram, representations of functional connectivity that depend on both frequency and time and are similar to a spectrograms, (Fig. 5A) were constructed during spindles using a complex Morlet wave convolution for 12-17 Hz using the “freqanalysis” function with the “wavelet” method from the Matlab Fieldtrip package. The wavelet cycle width was set to 7 with a length defined as 3 standard deviations (SDs) of the implicit Gaussian kernel. Connectivity analysis utilizing coherograms was performed using the “connectivityanalysis” function from the Matlab Fieldtrip package. Coherograms were averaged across specified channels over spindle occurrence times (500 ms preceding and 1500 ms following spindle start). Maximal imaginary coherence was identified over all coherogram time-frequency bins, 11 frequency components with 0.5 Hz steps from 12 to 17 Hz over 2000 time points for the 2000 ms was analysed, and used to quantify the number of time-frequency bins exceeding 70% of the maximum imaginary coherence.

**Figure 4.**
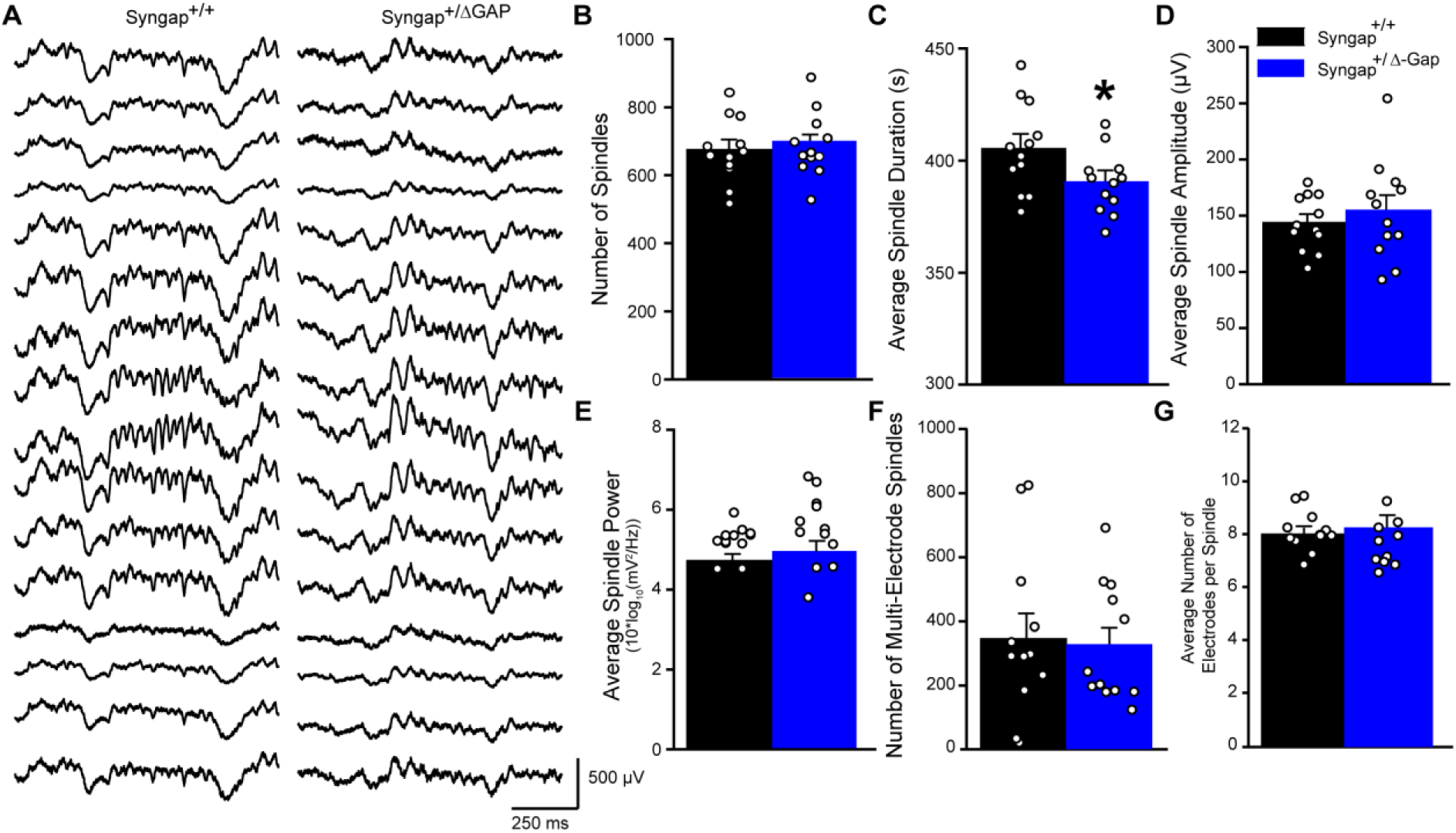
Sleep spindles are unaltered in Syngap^+/Δ-GAP^ animals. (A) Representative EEG traces during sleep spindles in 16 electrodes in Syngap^+/+^ (left) and Syngap^+/Δ-GAP^ (right) rats. Plots of spindle number (B), average duration (C), average amplitude (D), average power in the spindle frequency band (12 – 17 Hz) (E), multi-electrode spindles (F) and average electrodes per spindle (F). Bars indicate mean values (mean ± standard error of the mean (SEM)). Points correspond to values from individual rats. There was a significant decrease in spindle duration in Syngap^+/Δ-GAP^ rats with no differences in the other metrics (* = p<0.05, unpaired two-sample t-tests).

**Figure 5.**
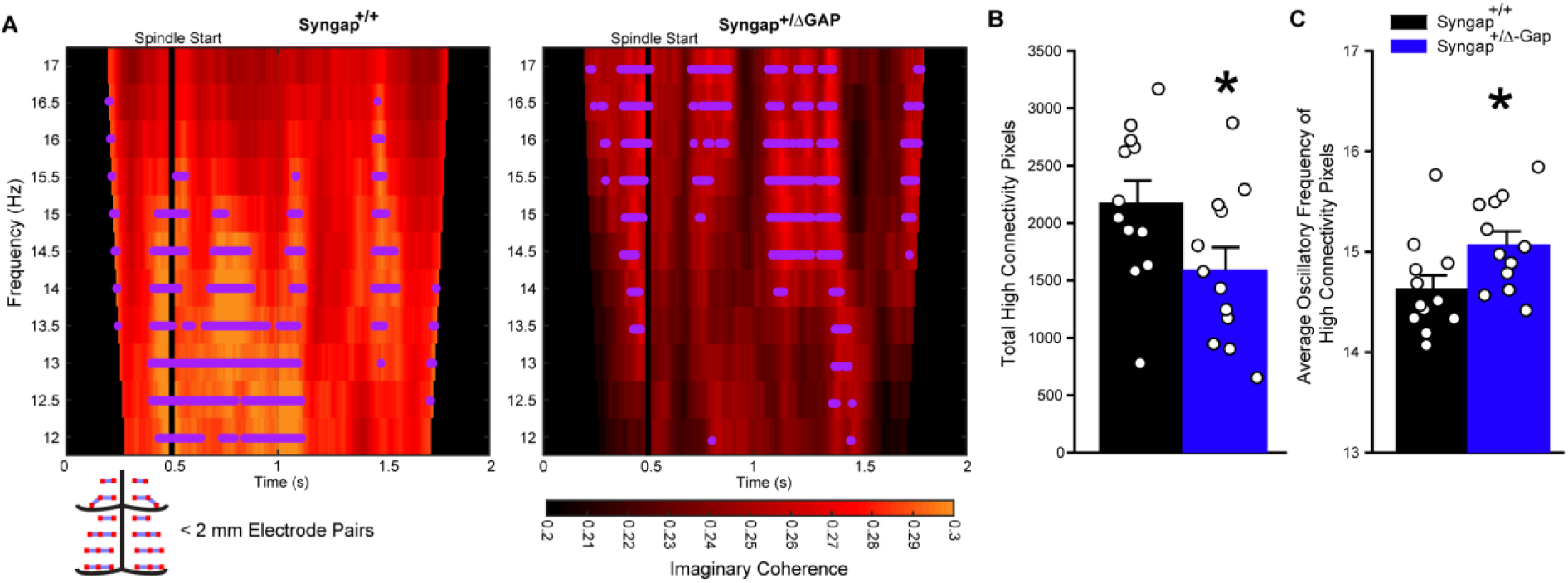
The occurrences of high connectivity during sleep spindles was reduced in short-distance (< 2 mm apart) electrode pairs in Syngap^+/Δ-GAP^ rats. (A) Average coherograms across electrode pairs preceding and during sleep spindles (top). Magenta dots indicate instances of high connectivity (70% of maximum average connectivity time-frequency bin detected in each animal). Schematic of electrode pairs < 2mm apart and imaginary coherence colour scale (below). Plots of total high connectivity time-frequency bins (B) and average oscillatory frequency of high connectivity pixels per spindle (F). Bars indicate mean values (mean ± SEM). Points correspond to values from individual rats. There was a significant decrease in high connectivity time-frequency bins and a significant increase in the average oscillatory frequency of high connectivity time-frequency bins in Syngap^+/Δ-GAP^ animals (* = p<0.05, unpaired two-sample t-tests).

### Statistics

When normality and homoscedasticity were above the rejection value of < 0.05 (estimated by Shapiro–Wilk and Levene’s tests respectively), between genotype comparisons across animals and electrodes were made with two-sample unpaired t-tests. Otherwise, a two-sample Mann Whitney-U rank sum test was used for unpaired two-sample comparison. A two-way ANOVA was utilized to compare the average Z’ imaginary coherence in commonly used frequency bands. We utilized a non-parametric two-sided permutation analysis with 40,000 simulation runs to compare average coherence per frequency band and over time during sleep spindles across electrode pairs. Consecutive significant frequencies clusters determined ranges with significant differences^23^.

In this study we analysed data recorded from *Syngap*^+/*Δ-GAP*^ and *Syngap*^+/*+*^, which previously allowed us to show significant differences in the incidence and duration of absence seizures^17^. For our a priori statistical power calculations for the present study we utilized alpha = 0.05. For comparisons between genotypes of percent time, total minutes and average duration of individual brain state bouts we estimated based on changes in brain states in mutant mice from similar results reported in another neurodevelopmental model^24^. For total minutes we set the mean of the control to 75 min, the mean of the mutant group to 55 min and the standard deviation for the mutants was 15.8. For n = 12 the resulting power is 0.803 and effect size 1.27.

## Data Availability

Upon article acceptance data will be made freely available on Edinburgh DataShare from the University of Edinburgh (https://datashare.ed.ac.uk/) and can be obtained directly from the corresponding author.

## Funding

The work performed here was funded by The Simons Initiative for the Developing Brain, the Patrick Wild Centre, an Epilepsy Research UK grant [F1603] and a Wellcome Trust Institutional Support Fund grant [204804/Z/16/Z]. For the purpose of open access, the author has applied a CC BY public copyright licence to any Author Accepted Manuscript version arising from this submission.

## Author Contributions

P.C.K., J.E., S.M.T. and A.G-S. designed the experiments. I.B.P. performed the experiments. I.B-P., J.M-R. and Z. L. analysed the data. A.G-S. wrote the manuscript with input from all authors.

## Results

### Sleep-wake distribution and spectral properties in *Syngap*^+/*Δ-GAP*^ rats

We previously reported that Syngap^+/Δ-GAP^ rats displayed absence seizures more frequently than littermate controls utilizing six-hour EEG (Fig. 1A, Supplementary Fig. 1) recordings when animals were in a quiet-wake state^17^. To determine whether *Syngap*^+/*Δ-GAP*^ animals have abnormalities in their brain state distribution, we classified all individual 5 s recording epochs from those previous recordings as NREM sleep, REM sleep or wake (Fig. 1A).

We found that *Syngap*^+/*Δ-GAP*^ rats spent an equivalent percentage of time in all states when compared to wild-type littermate controls (unpaired two-sample t-test for REM: DF = 22, T = 0.67, p=0.51; NREM: DF = 22, T = 0.0025, p = 0.1; wake: DF = 22, T = 0.26, p=0.8; Fig. 1B).

Nonetheless, *Syngap*^+/*Δ-GAP*^ animals had a significantly lower number of wake and NREM bouts, with REM bouts remaining unchanged (unpaired two-sample t-test for REM: DF = 22, T = 0.21, p=0.83; wake: DF = 22, T = 3.8, p= 0.00095; NREM sleep: DF = 22, T = =3.5, p=0.002; Fig. 1C). This coincided with increased average bout duration during wake and NREM and, no-difference in REM bout duration (unpaired two-sample t-test for REM: DF = 22, T = 0.79, p=0.44; wake: DF = 22, T = 2.57, p=0.017; two-sample Mann Whitney-U rank sum test for NREM: U=25, p=0.007, Fig. 1D). Therefore, during 6 hr recordings, *Syngap*^+/*Δ-GAP*^ rats display an abnormal brain state distribution.

We calculated the average EEG spectral power from one representative channel across all animals and epochs of each brain state and found no significant differences between *Syngap*^+/*Δ-GAP*^ rats and their wild-type littermates in commonly utilized frequency bands during NREM (unpaired two-sample t-test for delta: DF = 22, T = 0.05, p=0.96; theta: DF = 22, T = 1.8, p=0.08; low gamma DF = 22, T = 1.16, p= 0.26; two-sample Mann Whitney-U rank sum test for sigma: U= 41, p=0.08 Fig. 1E and H), REM (two-sample unpaired t-test for delta: DF = 22, T = 0.008, p=0.99; theta: DF = 22, T = 0.24, p=0.81; low gamma DF = 22, T = 0.28, p= 0.78; two-sample Mann Whitney-U rank sum test for sigma: U= 70, p=0.93, Fig. 1F and I) or wake (two-sample unpaired t-test for delta: DF = 22, T = 0.51, p=0.61; sigma: DF = 22, T = 0.95, p=0.35; low gamma DF = 22, T = 0.32, p= 0.76; two-sample Mann Whitney-U rank sum test for theta: U= 56.5, p=0.39, Fig. 1G and Fig. 1J). These data show that *Syngap*^+/*Δ-GAP*^ rats display an abnormal sleep-wake distribution, although overall spectral properties were unchanged.

### Functional connectivity in *Syngap*^+/*Δ-GAP*^ rats

Abnormalities in synaptic connectivity may underlie the cognitive pathologies and epilepsy in *SYNGAP* haploinsufficiency^25,26^. Network activity pathophysiology in neurodevelopmental disorders may be detected by analysing connectivity between EEG electrodes^27^. We therefore analysed the imaginary coherence, a generalization of correlation in the phase domain^21^, between voltage signals in our multi-site recordings during each brain state. Imaginary coherence decreases the likelihood of false dependencies between electrodes due to volume conductance by correlating between signal phases and not amplitude. We calculated the imaginary coherence for frequencies from 0 to 50 Hz for each brain state averaged across all epochs for all 496 combinations of electrode pairs from the 32 electrodes recorded. Since short or long distance connections may be differentially affected in neurodevelopmental disorders and may display varying levels of correlation^22^, imaginary coherence values were then averaged by electrodes that were grouped by short or long distances (Fig. 2, Fig. 3, Supplementary Fig. 1 and Supplementary Fig. 2).

As rodent surface multi-site EEG probes have not been previously utilized to compare short and long-distance electrode connectivity, we calculated whether there were differences between *Syngap*^+/*Δ-GAP*^ rats and wild-type controls using multiple distance thresholds separating groups of channels into short and long-distance channel combinations (Fig. 2, Fig. 3 and Supplementary Fig. 2). The most striking differences occurred during NREM amongst short-distance pairs < 2 mm from each other (Fig. 2A, Fig. 2B, Fig. 2C and Supplementary Fig. 2). When comparing commonly used frequency bands there was a clear, although non-significant, trend towards decreased theta and sigma band imaginary coherence in *Syngap*^+/*Δ-GAP*^ rats (two-way ANOVA, p = 0.06, F = 3.54, df = 1, Fig. 2A and Fig. 2B). To identify abnormal frequency bands, we utilized an unbiased approach in which we compared the imaginary coherence between the two groups by statistically testing each individual 0.5 Hz frequency bin between 0.5 and 50 Hz and plotting p-values as a function of frequency^23^. We found clusters of consecutive significantly different frequencies with p ≤ 0.05 corresponding to frequencies with significantly decreased *Syngap*^+/*Δ-GAP*^ imaginary coherence (two-sided randomization-based non-parametric test with cluster-based multiple comparison correction, Fig. 2C). These clusters occurred between 5 and 9 Hz and between 11.5 and 29.5 Hz. Furthermore, there was a striking subset of consecutive frequencies with lower p-values (p ≤ 0.01) between 12 and 22 Hz (Fig. 2C and Supplementary Fig. 2B). That range overlaps with the reported frequency range of sleep spindles (12 to 17 Hz), which are characteristic of NREM sleep and are thought to be critical for normal brain function and memory consolidation^28,29^. This cluster of significantly lower imaginary coherence frequencies suggests that connectivity may be compromised specifically during sleep spindles in *Syngap*^+/*Δ-GAP*^ animals in short-range distance electrode pairs.

There were no significant differences in commonly utilized frequency bands in short-distance electrodes < 2 mm during REM (two-way ANOVA, p = 0.75, F = 0.10, df = 1, Fig. 2D and Fig. 2E) although a trend towards significance in the sigma band was evident in wake periods (two-way ANOVA, p = 0.10, F = 2.72, df = 1, Fig. 2G, H). In contrast to NREM, there was only a small cluster of frequencies between 16 and 18 Hz during REM with p ≤ 0.05 (two-sided randomization-based non-parametric test with cluster-based multiple comparison correction, Fig. 2F). While in wake there were no significantly different frequencies (two-sided randomization-based non-parametric test with cluster-based multiple comparison correction, Fig. 2I).

We also compared electrode pairs > 2 mm from each other as long-distance connectivity has been shown to be increased in children with other neurodevelopmental disorders^22^. There were no significant differences in standard frequency bands between genotypes during NREM (two-way ANOVA, p = 0.57, F = 0.33, df = 1, Fig. 3A and Fig. 3B) or REM (two-way ANOVA, p = 0.76, F = 0.09, df = 1, Fig. 3E and Fig. 3F). This was confirmed by a lack of significantly different frequency clusters in both NREM and REM (two-sided randomization-based non-parametric test with cluster-based multiple comparison correction, Fig. 3C and Fig. 3F). However, a distinct non-significant trend towards increased theta and sigma coherence in *Syngap*^+/*Δ-GAP*^ rats during wake in long distance electrode pairs was detected (two-way ANOVA, p = 0.06, F = 3.72, df = 1, Fig. 3G and Fig. 3H). Clusters of consecutive significant frequencies, with p ≤ 0.05 showing higher connectivity in *Syngap*^+/*Δ-GAP*^ rats, were present during wake in the theta range between 8 and 9.5 Hz, and in the sigma range between 14 and 16.5 Hz (two-sided randomization-based non-parametric test with cluster-based multiple comparison correction, Fig. 3I), suggesting hyper-connectivity phenotypes may be present during wake in *Syngap*^+/*Δ-GAP*^ rats.

Overall, these data show that *Syngap*^+/*Δ-GAP*^ rats have connectivity abnormalities between electrode pairs located at short and long-distances from each other, with particularly striking deficiencies in imaginary coherence during NREM amongst short-distance pairs of electrodes.

### Sleep spindles in *Syngap*^+/*Δ-GAP*^ rats

Due to the reduction in average imaginary coherence in the sleep spindle frequency range in short-distance electrode combinations, we assessed whether sleep spindles were abnormal in *Syngap*^+/*Δ-GAP*^ rats (Fig. 4A). We first automatically detected spindles across animals from a single channel, over the right primary somatosensory cortex (Supplementary Fig. 1), to determine whether spindle number and duration is altered in *Syngap*^+/*Δ-GAP*^ rats. There was no significant difference in the total number of spindles detected (two-sample unpaired t-test, DF = 22, T = 0.59, p = 0.56, Fig. 4B) although there was a small decrease in the average duration of spindles in *Syngap*^+/*Δ-GAP*^ rats when compared to wild-type littermates (two-sample unpaired t-test, DF = 22, T = 2.19, p = 0.039, Fig. 4C). The average spindle amplitude was not significantly different between both genotypes (two-sample unpaired t-test, DF = 22, T = 0.75, p = 0.461, Fig. 4D) nor was the average power after spindle detection in the 12 to 17 Hz band (Mann-Whitney u-test, U = 52, p = 0.26, Fig. 4E). We then automatically detected spindles across all channels during NREM to assess whether there was a deficit between how spindles spread between short-distance pairs of electrodes. We found that there was no significant difference between *Syngap*^+/*Δ-GAP*^ rats and wild-type littermates in both the number of times that a spindle was simultaneously identified in more than one electrode (two-sample unpaired t-test, DF = 22, T = 1.78, p=0.26 Fig. 4F) and the number of electrodes in which a spindle was present when it was detected in more than one electrode (two-sample unpaired t-test, DF = 22, T = -0.92, p=0.37, Fig. 4G). These results show that spindle occurrence, amplitude, spectral power and detection of spindles throughout the cortex are unchanged in *Syngap*^+/*Δ-GAP*^ rats.

### Dynamic functional connectivity in *Syngap*^+/*Δ-GAP*^ rats during sleep spindles

We hypothesized that the significant reduction in average imaginary coherence in *Syngap*^+/*Δ-GAP*^ rats during NREM in short-distance electrode pairs may be a consequence of deficits in connectivity across channels during spindles. We therefore calculated average imaginary coherograms during spindles across all electrode combinations at distances ≤ 2 mm from each other for each animal, and we identified the total number of time-frequency bins in the coherograms that exceeded 70 % of the maximum connectivity detected in each individual animal (Fig. 5A). We found that *Syngap*^+/*Δ-GAP*^ rats had significantly decreased total high connectivity time-frequency bins from average coherograms when compared to wild-type controls (two-sample unpaired t-test, DF = 22, T = -2.39, p=0.03, Fig. 5B). This coincided with a significant increase in the spectral frequency at which high-connectivity time-frequency bins occurred (two-sample unpaired t-test, DF = 22, T = 2.13, p=0.04, Fig. 5C). These data show that there is a deficit in the occurrences of high functional connectivity in *Syngap*^+/*Δ-GAP*^ rats during spindles that may contribute to the overall decrease in imaginary coherence during NREM amongst short-distance electrode pairs.

We also compared the average values of imaginary coherence from coherograms during the entire length of spindles between the two genotypes and found that there was no significant reduction both across averages from all 496 channel combinations (two-sample unpaired t-test, DF = 22, T = 1.34, p = 0.19, Fig. 6B) and across averages from pairs of channels ≤ 2 mm from each other (two-sample unpaired t-test, DF = 22, T = 1.72, p = 0.10, Fig. 6B). This suggests that what contributes to decreased spindle coherence in short-distance channels is the reduction of occurrences of high-connectivity during spindles and not the overall average across the entire 12 to 17 Hz range and time preceding and following the start of spindles.

**Figure 6.**
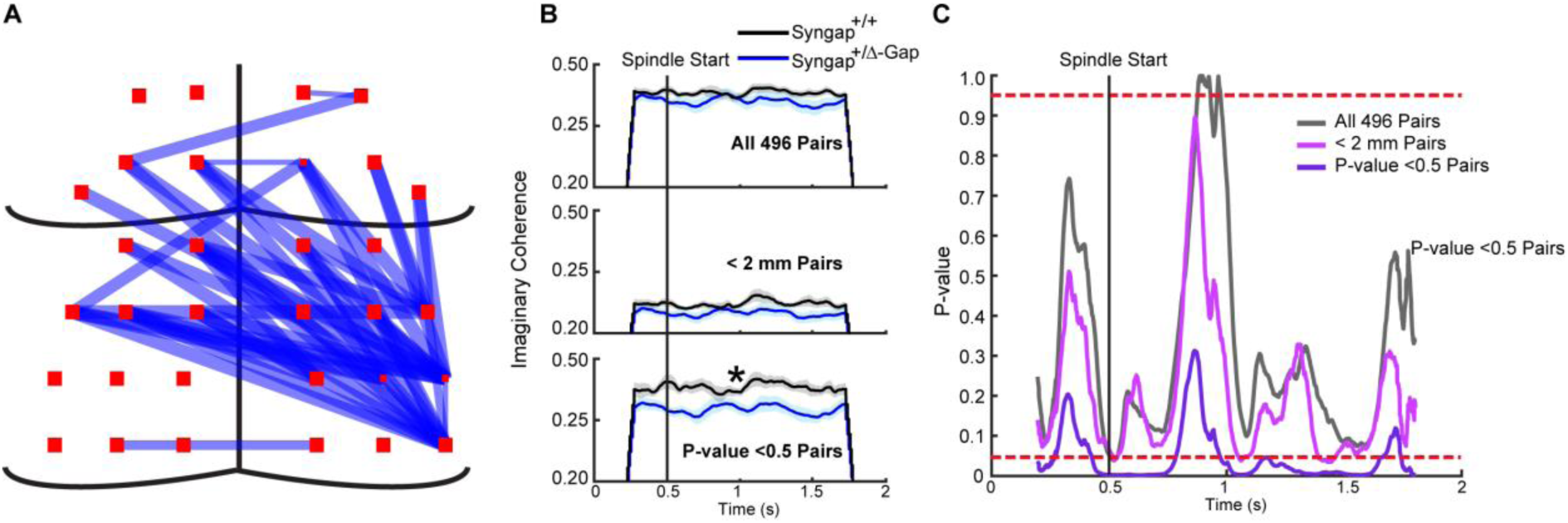
A specific subset of electrode pairs displays significantly reduced imaginary coherence during sleep spindles in Syngap^+/Δ-GAP^ rats. (A) Schematic of electrodes pairs with a significant reduction in imaginary coherence during sleep spindles in Syngap^+/Δ-GAP^ animals. Note: Thickness of lines indicate relative significance level detailed in Supp Table 1. (B) Average imaginary coherence preceding and during sleep spindles for all 496 (top), < 2 mm apart (middle) and individually significant (bottom) electrode pairs (Supp Table 1). Shaded area indicates SEM. (C) Plots of p-values for cluster-based nonparametric test preceding and during sleep spindles for all 496, < 2 mm apart and individually significant electrode pairs. Dotted red lines indicate two-sided p-value thresholds of ≥ 0.975 and ≤ 0.025 corresponding to significantly different thresholds equivalent to p ≤ 0.05.. Note: Clusters of significant times during sleep spindles were primarily found in individually significant electrode pairs in Syngap^+/Δ-GAP^ rats.

It is possible that specific channel combinations that are not organized by distance but by cortical location may have altered coherence during spindles. We therefore analysed whether specific channel pairs had significantly decreased connectivity when compared across animals and we found 45 combinations of channels where the imaginary coherence was significantly decreased in *Syngap*^+/*Δ-GAP*^ rats between 12 and 17 Hz (Fig. 6A and Supplementary Table 1). These channels were located caudal to bregma predominantly on the right posterior hemisphere over somatosensory, association and visual cortices (Fig. 6A and Supplementary Fig. 1). Some long-range and interhemispheric connections also showed differences between the groups (Fig. 6A). When we evaluated the average imaginary coherence for these channel combinations, we found a significant decrease in *Syngap*^+/*Δ-GAP*^ rats compared to controls (two-sample unpaired t-test, DF = 22, T = 3.04, p = 0.006, Fig. 6B). This was confirmed by comparing specific time points before and during spindles between the two genotypes utilizing the different combinations of electrode pairs. Clear and long clusters of consecutively significant timepoints, p ≤ 0.05, were present when utilizing the 45 channels with significantly decreased coherence and not when utilizing all 496 pairs or electrodes < 2 mm from each other (two-sided randomization-based non-parametric test with cluster-based multiple comparison correction, Fig. 6C).

Overall, these results show that there are deficits in instances of high functional connectivity during spindles amongst electrode combinations at short distances from each other, while a subset of electrode combinations have a higher degree of connectivity deficit suggesting potential deficits in *Syngap*^+/*Δ-GAP*^ rats in specific underlying cortical anatomy.

## Discussion

We show that brain states are altered in *Syngap*^+/*Δ-GAP*^ animals as the number of NREM and wake bouts are decreased. Nonetheless a corresponding increase in average NREM and wake bout duration in *Syngap*^+/*Δ-GAP*^ rats resulted in an equal percentage of time spent in NREM, REM and wake states when compared to wild type controls. Although no significant differences in spectral power or imaginary coherence were detected in commonly utilized frequency bands; when imagery coherence was compared between genotypes with a non-biased non-parametric method, clear decreases in *Syngap*^+/*Δ-GAP*^ animals in electrode pairs < 2 mm apart during NREM between 11.5 and 29.5 Hz were present. The overlap of the decrease in imaginary coherence during NREM with the sleep spindle frequency range amongst pairs of electrodes < 2 mm apart, prompted an in-depth analysis in sleep spindle properties. The occurrence, amplitude, power and detection of sleep spindles across multiple electrodes was unchanged in *Syngap*^+/*Δ-GAP*^ rats with only a small decrease in duration detected. Notwithstanding, dynamic coherence analysis revealed that during spindles, instances of high connectivity were decreased in *Syngap*^+/*Δ-GAP*^ animals, which was accompanied by an increase in average high-connectivity imaginary coherence frequency. Finally, the average coherence during spindles amongst all pairs of electrodes, as well as those < 2 mm apart, was not significantly different between genotypes despite the decrease in the instances of high connectivity for electrodes < 2mm in *Syngap*^+/*Δ-GAP*^ rats. However, by identifying a subset of electrodes with significantly decreased connectivity during spindles in *Syngap*^+/*Δ-GAP*^ animals, we found that as a group, these electrodes did show average decreased connectivity. These electrodes were primarily located in the right caudal hemisphere, suggesting that cortical dynamics at this location may be particularly affected by the *SYNGAP1* mutation.

Difficulties in both initiating and maintaining sleep as well as reduced overall sleep duration have been reported in individuals with *SYNGAP1* mutations^3,8,9^. In our previous work we performed 6 hr recordings EEG recordings and found that *Syngap*^+/*Δ-GAP*^ animals displayed absence seizures at a higher rate than controls ^17^. We have further analysed these recordings, which were made during daylight hours, resulting in rats spending approximately half of the recording time asleep. Despite recent clinical reports, we found that the overall time spent asleep was not different from littermate controls in mutant animals. Nonetheless, the number of times that *Syngap*^+/*Δ-GAP*^ rats entered NREM or wake brain states was decreased. Interestingly, there was also an increase in the duration of NREM and wake bouts, which suggest that overall sleep structure is compromised in these animals. Since sleep is intermingled with wake states throughout the day in rats, full-day circadian recordings may identify further sleep abnormalities in this mutant line.

EEG has been suggested as a potential method to identify clinical biomarkers in NDDs ^30^ and indeed analysis of EEG connectivity and networks have yielded specific signatures for epilepsy and autism^22,31^. Here, we performed analysis on the connectivity between electrode pairs from our multi-site EEG probes. We utilized imaginary coherence since the likelihood of volume conductance artefacts with a method analysing amplitude correlations, may be higher with the reduced skull size of rats compared to humans.

We separated electrode pairs between short and long distance groups and averaged the overall coherence within these groups. We performed comparisons with varying threshold distances separating short and long distance electrode pair groups and found that the clearest deficits in connectivity in *Syngap*^+/*Δ-GAP*^ rats occurred during NREM in short distance electrodes < 2 mm. SYNGAP is primarily located in excitatory synapses and is a key regulator of spine formation^14–16^. Cortical pyramidal cells form most of their intracortical connections with neighbours in close proximity^32^. These data suggest that there may be a deficit in these connections, which may be heightened during cortical activity in NREM.

One of the characteristics of NREM are sleep spindles, which are thought to be critical for memory processes^29^, and lower imaginary coherence in the spindle frequency range led us to investigate their characteristics in *Syngap*^+/*Δ-GAP*^ animals. We found that their overall characteristics were unaffected, as well as the average connectivity during a spindle. Nonetheless, brief periods of high connectivity were significantly decreased in *Syngap*^+/*Δ-GAP*^ rats, with a concurrent increase in the oscillatory frequency at which these high connectivity events occurred. How cortical neuronal populations interact during sleep spindles may be compromised in *Syngap*^+/*Δ-GAP*^ animals. Analysis of sleep spindle connectivity between individual channel combinations showed that a subgroup of connections had significantly lower imaginary coherence in *Syngap*^+/*Δ-GAP*^ rats. This area is located over the right somatosensory, association and visual cortices, which may be more vulnerable to deficits due to *SYNGAP1* mutation. Some interhemispheric connectivity abnormalities may also be present, although the overall average imaginary coherence between long-distance connections was not significantly altered during NREM (Fig. 3C).

Finally, we saw that despite decreases in the overall number of NREM and wake bouts, the overall time spent in those states was not decreased due to their increased duration in *Syngap*^+/*Δ-GAP*^ animals. Interestingly, we also saw decreased connectivity during NREM in the sigma range in mutant animals, which was accompanied by increased connectivity in the theta and sigma range during wake. A degree of homeostatic adaptation may be occurring in mutant animals to counteract the effects of the mutation during different brain states.

In conclusion, we report sleep abnormalities and differences in connectivity during specific brain states in the *Syngap*^+/*Δ-GAP*^ rat model. Imaginary coherence analysis of EEG data may have value as a clinical biomarker and this analysis points to specific neuronal populations that may be affected by the mutation.

