## Supplementary Figures and Tables for "Abnormal brain state distribution and network connectivity in a *SYNGAP1* rat model"

**Supplementary Material**

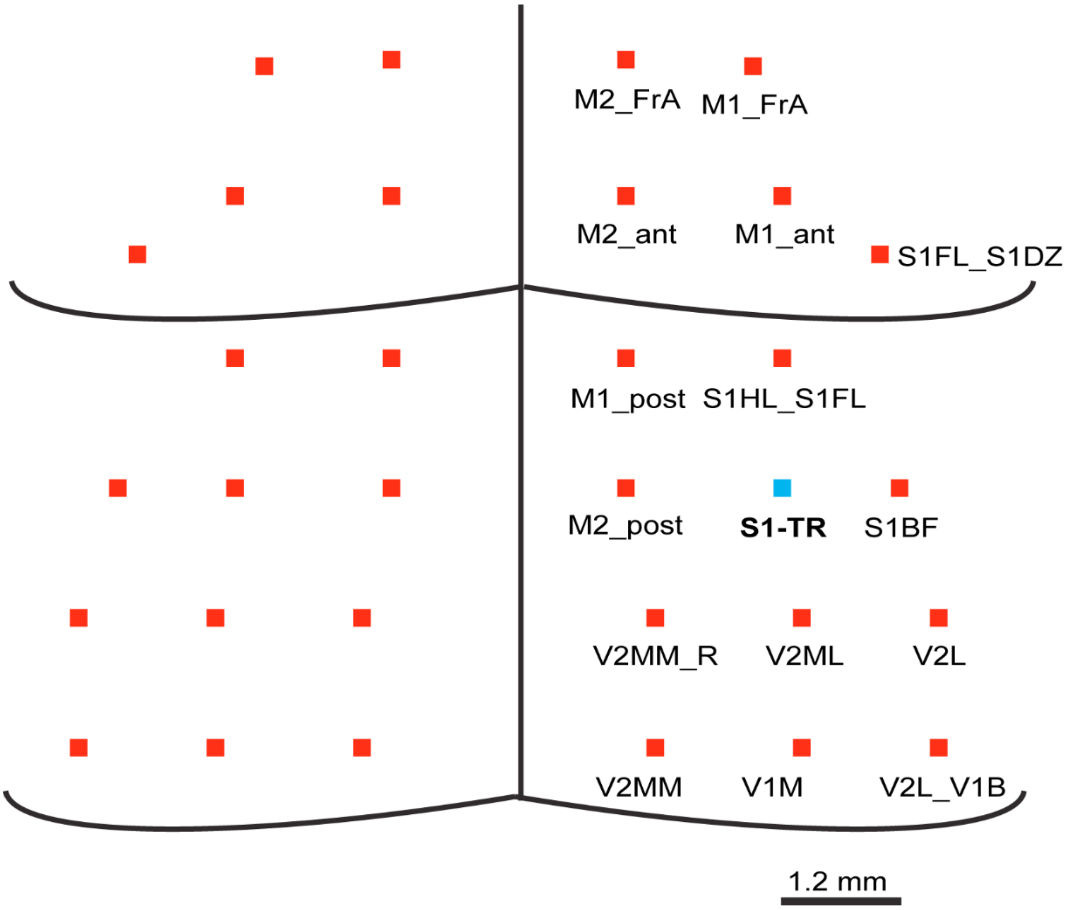


**Supp Fig 1. Diagram of approximate electrode location and abbreviations of approximate underlying cortical areas.**
Coordinates were based on approximate locations from (Paxinos and Watson, 1998)

V2L_V1B: Secondary visual cortex lateral area – Primary visual cortex binocular area
V1M: Primary visual cortex monocular area
V2MM: Secondary visual cortex mediomedial area
V2L: Secondary visual lateral area
V2ML: Secondary visual mediolateral area
V2MM_R: Secondary visual cortex mediomedial area - retrosplenial agranular area
S1DZ_S1BF: Primary somatosensory cortex barrel Field
S1Tr: Primary somatosensory cortex trunk area
M2_post: Posterior secondary motor cortex
S1HL_S1FL: Primary somatosensory cortex hindlimb region – forelimb region
M1_post: Posterior primary motor cortex
S1FL_S1DZ: Primary somatosensory cortex forelimb region - disgranular region
M1_ant: Anterior primary motor cortex
M2_ant: Anterior secondary motor cortex
M1_FrA: Primary motor cortex – frontal association cortex
M2_FrA: Secondary motor cortex – frontal association cortex

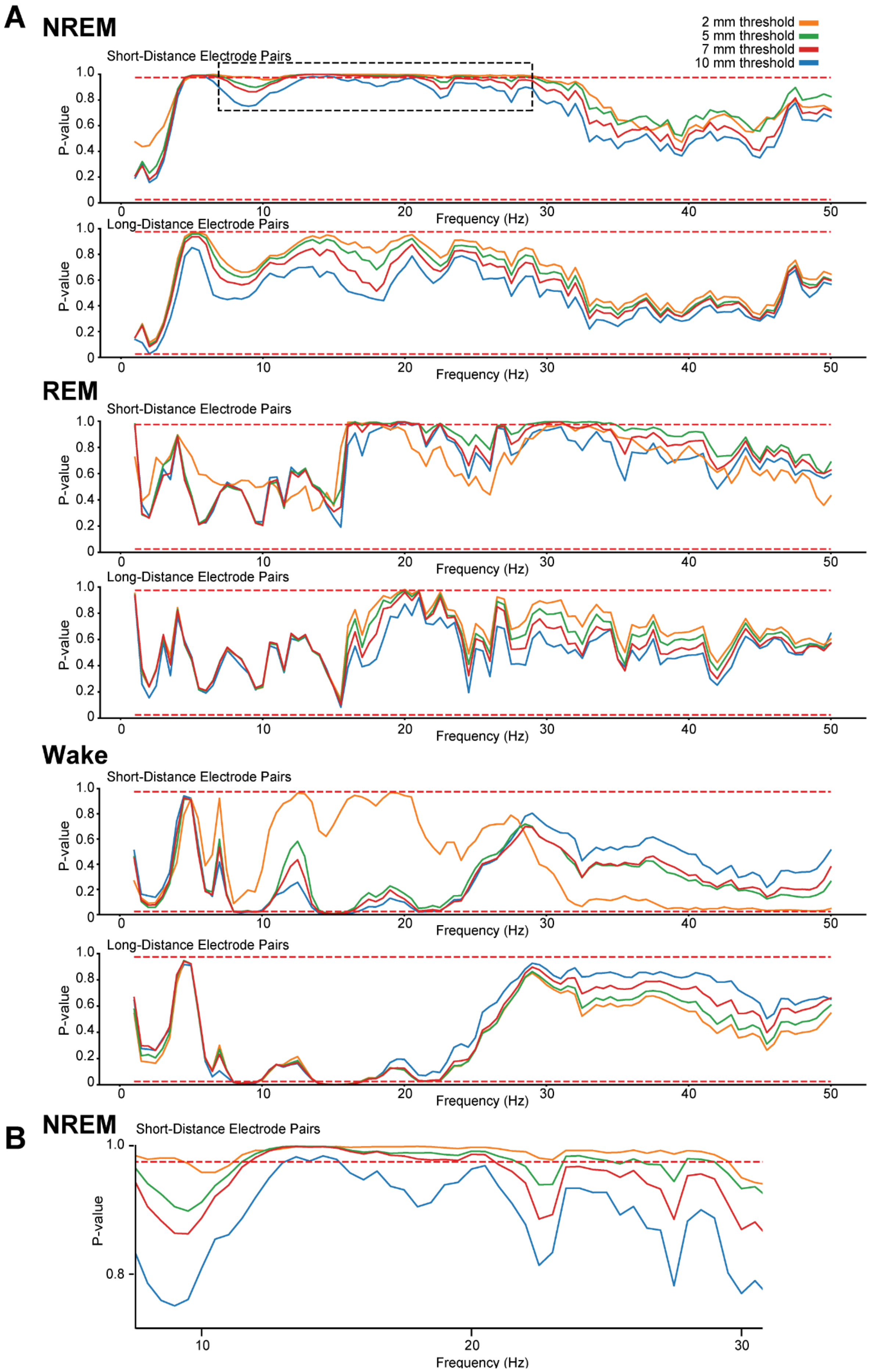
A
NREM
2 mm threshold
5 mm threshold
7 mm threshold
10 mm threshold
Short-Distance Electrode Pairs
Long-Distance Electrode Pairs
P-value
Frequency (Hz)
REM
Short-Distance Electrode Pairs
Long-Distance Electrode Pairs
Wake
Short-Distance Electrode Pairs
Long-Distance Electrode Pairs
B
NREM
Short-Distance Electrode Pairs
P-value
Frequency (Hz)

**Supp Fig. 2. Plots of p-values at multiple distance thresholds.** (A) P-values of differences at individual frequencies comparing imaginary coherence between *Syngap*$^{+/\Delta\text{-}GAP}$ and *Syngap*$^{+/+}$ rats for cluster-based nonparametric tests for short and long distance electrodes during NREM, REM and wake. Dotted red lines indicate two-sided p-vale thresholds of $\geq 0.975$ and $\leq 0.025$ corresponding to significantly different thresholds equivalent to $p \leq 0.05$. Note: The distance threshold determines how electrode combinations are grouped as short or long distance combinations. At thresholds of 2, 5, 7 and 10 mm there were 20, 170, 284 and 432 electrodes in the short distance groups and 476, 326, 212 and 64 electrodes in the long distance groups respectively. (B) Expanded p-value and frequency plot of area within the dotted black rectangle in (A). Note: A long cluster of significant frequencies was found during NREM between 11.5 and 29.5 Hz in electrodes ≤ 2mm indicating a decrease in Z' imaginary coherence in *Syngap*$^{+/\Delta\text{-}GAP}$ rats. The longest consecutively significant frequencies clusters are present amongst short-distance electrodes ≤ 2mm apart.

**Supp Table 1. Electrode pairs during sleep spindles with significantly decreased dynamic imaginary coherence.** All 496 electrode pairs were compared across animals with a two-sample t-test. The 45 significant pairs are listed below with corresponding p-values.

| Electrode Pairs | | p-values |
|---|---|---|
| V2L_V1B_RIGHT | S1Tr_LEFT | 0.031 |
| V2L_RIGHT | S1Tr_LEFT | 0.033 |
| V2ML_RIGHT | S1Tr_LEFT | 0.031 |
| V2MM_RSA_RIGHT | S1Tr_LEFT | 0.049 |
| M2_ant_RIGHT | S1DZ_S1BF_LEFT | 0.042 |
| V2L_V1B_RIGHT | S1DZ_S1BF_LEFT | 0.003 |
| V2L_RIGHT | S1DZ_S1BF_LEFT | 0.007 |
| V2ML_RIGHT | S1DZ_S1BF_LEFT | 0.050 |
| S1DZ_S1BF_RIGHT | S1DZ_S1BF_LEFT | 0.035 |
| V2MM_RIGHT | V1M_LEFT | 0.035 |
| V2L_V1B_RIGHT | M2_post_LEFT | 0.028 |
| V2L_RIGHT | M2_post_LEFT | 0.038 |
| V2MM_RSA_RIGHT | M2_post_LEFT | 0.028 |
| V2L_V1B_RIGHT | S1HL_S1FL_LEFT | 0.050 |
| V2L_RIGHT | S1HL_S1FL_LEFT | 0.043 |
| V2L_V1B_RIGHT | M1_post_LEFT | 0.017 |
| V2L_RIGHT | M1_post_LEFT | 0.017 |
| V2ML_RIGHT | M1_post_LEFT | 0.031 |
| V2MM_RSA_RIGHT | M1_post_LEFT | 0.040 |
| S1DZ_S1BF_RIGHT | M1_post_LEFT | 0.029 |
| V2L_RIGHT | S1FL_S1DZ_LEFT | 0.049 |
| M1_FrA_RIGHT | M1_ant_LEFT | 0.033 |
| V2L_V1B_RIGHT | M1_ant_LEFT | 0.036 |
| V2L_RIGHT | M1_ant_LEFT | 0.041 |
| M2_ant_RIGHT | M2_ant_LEFT | 0.029 |
| V2L_V1B_RIGHT | M2_ant_LEFT | 0.021 |
| V2L_RIGHT | M2_ant_LEFT | 0.022 |
| S1DZ_S1BF_RIGHT | M2_ant_LEFT | 0.037 |
| M1_FrA_RIGHT | M2_FrA_RIGHT | 0.046 |
| V2L_V1B_RIGHT | M2_ant_RIGHT | 0.018 |
| V2L_RIGHT | M2_ant_RIGHT | 0.020 |
| S1DZ_S1BF_RIGHT | M2_ant_RIGHT | 0.020 |
| V2L_RIGHT | M1_ant_RIGHT | 0.026 |
| S1DZ_S1BF_RIGHT | M1_ant_RIGHT | 0.042 |
| V2L_RIGHT | S1FL_S1DZ_RIGHT | 0.025 |
| S1DZ_S1BF_RIGHT | S1FL_S1DZ_RIGHT | 0.044 |
| V2L_V1B_RIGHT | M1_post_RIGHT | 0.015 |
| V2L_RIGHT | M1_post_RIGHT | 0.008 |
| V2ML_RIGHT | M1_post_RIGHT | 0.037 |
| S1DZ_S1BF_RIGHT | M1_post_RIGHT | 0.004 |
| V2L_RIGHT | V2L_V1B_RIGHT | 0.002 |
| V2ML_RIGHT | V2L_V1B_RIGHT | 0.038 |
| S1DZ_S1BF_RIGHT | V2L_V1B_RIGHT | 0.008 |
| S1Tr_RIGHT | V2L_V1B_RIGHT | 0.023 |
| S1Tr_RIGHT | V2L_RIGHT | 0.020 |
